# Umbilical-brain endothelial communication via TSP-1 is linked with reduced brain angiogenesis in offspring of preeclampsia

**DOI:** 10.64898/2026.03.19.713060

**Authors:** Esthefanny Escudero-Guevara, Felipe Troncoso, Hermes Sandoval, Cristian Vargas, Marcelo Alarcón, Hiten D. Mistry, Lesia O. Kurlak, Rodrigo Moore-Carrasco, Jesenia Acurio, Carlos Escudero

## Abstract

**Background:** Preeclampsia, a maternal hypertensive syndrome affect fetal brain development and cerebral angiogenesis, with potential acute and long-term consequences. Underlying mechanisms of these brain vascular alterations are unknown. This study investigates the role of thrombospondin-1 (TSP-1), an antiangiogenic glycoprotein, as a key mediator of communication between the fetoplacental and fetal brain endothelium in the context of preeclampsia.

**Methods:** Conditioned media (CM) of human umbilical vein endothelial cells (HUVECs) from normal pregnancies (NP-CM) and preeclamptic pregnancies (PE-CM), were used to treat human (hCMEC/D3) and murine brain microvascular endothelial cells (BMECs). A proteomic analysis was performed in plasma of the umbilical cord of normal pregnancy and preeclampsia. TSP-1 was identify using proteomic analysis and confirmed by Western blot. PE-CM depleted of TSP-1, using immunoprecipitation, was used to evaluate protein-protein interaction with vascular endothelial growth factor (VEGF). Antibody-mediated blockage of TSP-1 was used to investigate antiangiogenic effect and pro-angiogenic signaling pathways in brain endothelial cells exposed to PE-CM.

**Results:** PE-CM significantly reduced angiogenesis, migration, and invasion of brain endothelial cells and altered cytoskeletal organization. These effects were accompanied by reduced VEGFR2 and AKT signaling, indicating impaired angiogenic pathways. Proteomic analysis of umbilical cord plasma revealed elevated TSP-1 levels in preeclampsia, which was confirmed by Western blotting. TSP-1 was also increased in PE-CM, and immunoprecipitation assays suggested a protein–protein interaction with VEGF. Antibody-mediated blockade of TSP-1 restored angiogenesis, as reflected by increased total tube length, and rescued VEGFR2 and AKT signaling in brain endothelial cells exposed to PE-CM.

**Conclusion:** TSP-1-mediated endothelium-endothelium communication between placenta-brain axis in offspring of mothers with preeclampsia. This communication mediated by TSP-1 may contribute to acute and long-lasting cerebrovascular dysfunction observed in infants exposed to preeclampsia.

## BACKGROUND

Preeclampsia is a hypertensive disorder of pregnancy accompanied by markers of multisystem organ damage affecting the kidneys, liver, or the brain ^1^. Preeclampsia affects approximately 2–8% of pregnancies worldwide and is the leading cause of maternal and fetal morbidity and mortality, especially in low-income countries ^2–4^.

While the maternal cardiovascular and neurocognitive sequelae has been well-documented in epidemiological studies ^5^, emerging evidence highlight a profound “vascular imprint” on the offspring. Infants born to women with preeclampsia face a 2- to 5-fold increased risk of perinatal stroke ^6–8^ or a 6-fold increased risk of neonatal encephalopathy. Furthermore, infants exposed to maternal hypertension during pregnancy, showed a higher incidence of subarachnoid hemorrhage ^9^. These clinical observations are underscored by case-control study in children born to women with preeclampsia ^10^ and experimental evidence in animal models ^11–13^ reporting cerebrovascular rarefactions and impaired neurovascular unit coupling. However, despite the gravity of these high neurovascular risk, the underlying mechanisms remain poorly understood.

The pathophysiology of preeclampsia and its perinatal consequences are rooted in placental insufficiency and the subsequent release of a bioactive “secretome” into both maternal and fetal compartments ^14–20^. For instance, elevated levels of the anti-angiogenic protein, sFlt-1, a decoy receptor of the vascular endothelial growth factor (VEGF), have been reported in umbilical cord blood from preeclamptic pregnancies ^21,22^. Nevertheless, the cellular origin of these factors within the fetal compartment and their functional consequences for fetal vascular development remain poorly comprehended. Crucially, the fetoplacental endothelium itself is now recognized as a potent source of pro-inflammatory and anti-angiogenic mediators ^23^.

Placental angiogenic imbalance directly affects fetal organ development. Thus, placental-targeted delivery of VEGF has been shown to improve fetal growth and survival in a mouse model of preeclampsia ^24^. Moreover, adenoviral-mediated placental gene therapy with VEGF (Ad.VEGF-A165) normalizes brain weight and prevents brain injury in offspring from a guinea pig model of fetal growth restriction, another condition characterized by placental insufficiency ^25^. These recent findings provide a “proof-of-concept” that restoring angiogenic equilibrium in the umbilical-fetal niche may mitigate neurodevelopmental injury.

Thrombospondin-1 (TSP-1) is a matricellular glycoprotein originally identified in platelets ^26^, and it is a well-established endogenous inhibitor of angiogenesis and growth factor signaling ^27,28^. Although TSP-1 has been investigated in maternal samples from women with preeclampsia showing conflicting results ^29–32^ its levels in the fetal circulation, particularly in umbilical cord blood, have not been estimated. Moreover, its potential role in fetal cerebral vascular development remains unexplored. Given its potent anti-angiogenic properties, we hypothesize that increased fetal exposure to TSP-1 may impair cerebral angiogenesis and contribute to the heightened cerebrovascular risk observed in offspring born to women with preeclampsia.

Therefore, we aimed to investigate whether TSP-1 released by umbilical vein endothelial cells reduces the angiogenic capacity of brain endothelial cells in the context of preeclampsia. We hypothesized that the fetoplacental endothelial secretome in preeclampsia contains elevated TSP-1, which impairs angiogenic signaling in brain endothelium.

## METHODS

### Human samples

Umbilical cord samples were obtained at the time of delivery from women with normotensive pregnancies (n=25) and from women diagnosed with preeclampsia (n=26). Preeclampsia was defined as new-onset hypertension after 20 weeks of gestation (systolic blood pressure ≥140 mmHg and/or diastolic blood pressure ≥90 mmHg on two separate measurements) accompanied by proteinuria >300 mg/L. Control pregnancies were normotensive and clinically uncomplicated. Study groups were matched for maternal age, gestational age at delivery, and newborn sex.

Fetal venous cord blood was collected immediately after delivery into EDTA-containing tubes at Nottingham University Hospitals (UK). Plasma was isolated by centrifugation, aliquoted, and stored at −80 °C until analysis. Clinical data, including maternal body mass index, parity, obstetric history, and birthweight percentile adjusted for gestational age, were recorded, as described previously ^11^.

All procedures conformed to the Declaration of Helsinki and were approved by the Ethics and Scientific Committee of Hospital Clínico Herminda Martín (Chillán, Chile, REF: Fondecyt 1240295- CEC-HCHM 806276-2024) and the HRA Research Ethics Committee, University of Nottingham (REF: 15/EM/0523). Written informed consent was obtained from all participants. Plasma samples were transported under controlled conditions to the Vascular Physiology Laboratory, Universidad del Bío-Bío, for endothelial functional studies.

### Animals

All animal procedures were approved by the Bioethics and Biosafety Committee of the University of Bío-Bío and conducted in accordance with the Guide for the Care and Use of Laboratory Animals and the principles of the 3Rs (Replacement, Reduction, and Refinement)^33^. Experiments are reported in compliance with ARRIVE 2.0 guidelines ^34^.

Female C57BL/6 mice (aged 2–4 months), were maintained under controlled environmental conditions (12:12 h light-dark cycle, temperature and humidity conditions, 25 °C) at the University of Bío-Bío Vivarium. Animals were fed a standard chow diet (Prolab RMH 3000, LabDiet, St. Louis, MO, USA) and had free access to water, as we previously described ^11,35^ Age-matched males were used to timed breeding. The presence of vaginal plug was designated as embryonic day 0.5 (E0.5).

### RUPP preeclampsia model

Pregnant females were randomly assigned to either the reduced uteroplacental perfusion pressure (RUPP) or sham group. On the gestational day 12.5 (E12.5), mice were anesthetized with inhaled isoflurane (induction 4%, maintenance 2%). A 5-mm midline abdominal incision was made, and both uterine horns were exteriorized. Ovarian and uterine arteries were bilaterally cauterized at four sites using low-temperature electrocautery (BoVie®, Florida, USA; 730 °C) until a 40–50% reduction in uteroplacental artery diameter was achieved. The uterus was returned to the abdominal cavity and the incision was closed in two layers using absorbable sutures. Sham-operated animals underwent identical surgical procedures without arterial cauterization.

This intervention produces placental ischemia and maternal features consistent with experimental preeclampsia, including impaired uteroplacental blood flow, as established in previous validation studies ^12,36,37^.

### Primary human umbilical vein endothelial cell (HUVEC) culture

Umbilical cord samples were washed with phosphate-buffered saline (PBS, pH 7.4), and both ends of the umbilical vein were cannulated. Endothelial cells were detached by incubation with collagenase type I (0.2 mg/mL, Gibco™, Waltham, MA, USA) for 10 min at 37 °C. The collagenase solution was collected and centrifuged (3500 rpm for 10 min). The resulting cell pellet was resuspended and seeded onto collagen–coated culture plates (Sigma Aldrich, MO, USA) in Medium 199 (Thermo Fisher Scientific, Waltham, MA, USA). Cells were maintained at 37 °C in a humidified incubator with 5% CO₂. The culture medium was changed every two days until 70–80% confluence was reached.

### Conditional media

HUVECs derived from normal pregnancies and pregnancies complicated by preeclampsia were cultured in Medium 199 (Thermo Fisher Scientific, Waltham, MA, USA) supplemented with 10% fetal bovine serum (FBS). Upon reaching ∼90% confluence, cells were washed with phosphate-buffered saline (PBS) and incubated in serum-free medium for 2 hours. Subsequently, cells were maintained in Medium 199 supplemented with 1% FBS for 48 hours to generate conditioned media (CM).

CM was collected and centrifuged at 3,500 rpm for 10 minutes to remove cellular debris, then stored at −80°C until further use. Experimental treatments were performed using CM derived from HUVECs of normal pregnancies (NP-CM) or preeclamptic pregnancies (PE-CM), normalized to 100 μg total protein, and applied at time points specified for each experimental protocol.

### Human brain endothelial cells

The human cerebral microvascular endothelial cell line, hCMEC/D3, was used for angiogenic experiments, as we previously described ^13^. Cells were cultured in EndoGRO™-MV Complete Culture Media Kit (Millipore), supplemented with 10% fetal bovine serum (FBS) and 1% penicillin–streptomycin. According to the manufacturer’s instructions, the medium contains endothelial growth supplements, including VEGF, fibroblast growth factor-2 (FGF-2), epidermal growth factor (EGF), insulin-like growth factor-1 (IGF-1), hydrocortisone, ascorbic acid, and heparin.

Cells were seeded on plates coated with type I collagen at a concentration of 50 µg/mL to support endothelial adhesion and phenotype. Cells were maintained at 37°C in a humidified atmosphere with 5% CO₂ and used between passages 8–12.

### Primary culture of brain endothelial cells

Isolation of murine brain endothelial cells (BMEC) were performed following Ruck *et al.,* protocol ^38^ with minor changes. Briefly, brains were obtained from E18.5 fetuses derived from sham and RUPP dams. Brainstems, cerebellum, thalamus, and meninges were removed. Tissue was digested using collagenase type II (1 mg/mL) and DNase I (0.1 mg/mL) for 1 h at 37 °C.

Cells were centrifuged and purified using a 20% BSA gradient, and then cultured until confluence. To isolate endothelial cells we used Dynabeads™ CD31 Endothelial Cell (Invitrogen™, Thermo Fisher Scientific, Waltham, MA, USA) following manufacturer’s protocol. Purified CD31 positive cells (CD31+) were cultured in DMEM supplemented with 20% FBS until experimental use.

### Murine brain endothelial cell line

The mouse brain endothelial cell line bEnd.3 was used in selected experiments. Cells were cultured in DMEM supplemented with 2 mM L-glutamine, 1 mM sodium pyruvate, 1× non-essential amino acids, 50 µM 2-mercaptoethanol, 10% FBS, and 1% penicillin–streptomycin. Cells were maintained on type I collagen-coated culture plates (Corning, 50 µg/mL).

For all endothelial models, cells were serum- and growth factor–deprived overnight prior to stimulation to minimize basal signaling activity.

### Cell viability assay

Viability tests were performed using the non-radioactive CellTiter 96® AQueous One Solution Cell Proliferation Assay (Promega, Madison, WI, USA). hCMECs/D3 were treated (6 h, 100 μg protein) with NP-CM or PE-CM, respectively. Absorbance was analyzed using an Epoch™ Microplate Spectrophotometer (BioTek Instruments, Winooski, VT, USA), with an absorbance measurement at 570 nm, as previously described ^39^.

### *In vitro* angiogenesis

The hCMECs/D3 or BMECs were seeded in a 96-well plate (30,000 cells per well) pre-coated with 60 μL of Matrigel® Basement Membrane Matrix (Corning Incorporated, Corning, NY, USA). The cells were treated with NP-CM or PE-CM, respectively (6 h, 100 μg protein, v/v). The angiogenesis *in vitro* was photographed using an inverted-phase contrast microscope at 10× magnification (Olympus, Tokyo, Japan). Network formation (branching, number of junctions, and total tube length) was quantified using the “Angiogenesis Analyzer” add-on of ImageJ v.5.2 software (NIH, USA), as previously described ^40,41^

### Cell migration

We use the “healing assay” on confluent hCMEC/D3. The cells were deprived of serum for 8 hours and then stimulated (6 h, 100 μg protein, v/v) with NP-CM or PE-CM, as described above. SPLScar™ Scratcher (SPL Life Sciences, Korea) was subsequently lined vertically in each of the wells. The event was recorded taking photos in microscopic fields (Olympus, Tokyo, Japan) and using an MShot MD90 camera (MShot Technology, GZ, China) every 3 hours for 12 hours. Images were quantified using Image J v5.2 software (NIH, USA). The percentage of migration between the initial area (T=0) and the migrated area at different times (T=3h, T=6h) was calculated using the following formula:

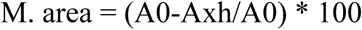

While the area M. represented the migratory area, A0 represented the area at time 0, or the bare area, and Axh represented the area that remained bare after 3 or 6 hours.

### *In vitro* invasion assay

We used transwell plates (Corning; polycarbonate membrane, 8 μm pore and 6.5 mm insert diameter) and Matrigel® Basement Membrane Matrix (Corning Incorporated, Corning, NY, USA) as the 3D matrix. Briefly, 60 μL of Matrigel was added to the upper chamber of the Transwell and incubated at 37 °C overnight. hCMEC/D3 were resuspended in a serum-free medium and seeded on Matrigel-coated membranes. 600 μL of conditioned media (NP-CM or PE-CM) were placed in the lower chambers. Transwell plates were incubated at 37°C and 5% CO₂ for 12 hours. The invaded cells were then fixed with 4% paraformaldehyde (PFA) and stained with crystal violet. The Matrigel was removed inside the insert, and the porous membrane with the invading cells was mounted on a microscope slide. Using an inverted-phase contrast microscope (Olympus, Tokyo, Japan), five representative microscopic fields were captured and the number of invading cells was quantified. As a positive control, a complete culture media supplemented with 10% FBS was used, given its known chemotactic effect. In contrast, baseline media without FBS was used as a negative control to establish the baseline level of invasion in the absence of chemoattractant stimuli.

### *Ex vivo* invasion test

For the *ex vivo* invasion assay, hCMEC/D3 were labeled with green fluorescent protein (GFP) by transduction with CellLight™ Nucleus-GFP, BacMam 2.0 (Thermo Fisher Scientific, Waltham, MA, USA). Subsequently, cerebral cortex sections were obtained from pups in embryonic stage E18.5, which were placed in the upper chamber of Transwell® inserts for 24 h. During this period, the lower chamber was maintained with culture medium supplemented with 10% FBS. After that, the experiment was initiated by carefully placing 1 × 10³ hCMEC/D3-GFP cells on the surface of the brain explant (putamen region). Next, 800 μL of either NP-CM or PE-CM were added in the lower chamber of the system. Positive and negative controls were also included. After 24 h of incubation, the brain explants were collected and fixed in 4% PFA. Subsequently, multiple serial sections Leica RM2045 microtome (Leica Microsystems Inc., Buffalo Grove, IL, USA) of the brain tissue were used to detect invasive hCMEC/D3-GFP cells by fluorescence microscopy (Motic BA410, Motic, Hong Kong, China).

### F-actin Cytoskeleton Analysis

F-actin organization was assessed in hCMEC/D3 cells following exposure to NP-CM or PE-CM (6 h, 100 µg total protein). Cells were fixed and stained with fluorescent phalloidin iFluor™ 488 (Abcam, Cambridge, UK; 1:500 dilution) to visualize filamentous actin and counterstained with DAPI (Invitrogen™, Thermo Fisher Scientific, Waltham, MA, USA) for nuclear identification. Samples were washed in PBS and mounted for fluorescence microscopy.

Images were acquired under identical settings using a fluorescence microscope (100× magnification). F-actin structures were analyzed using ImageJ v 5.5 software (NIH, USA) after spatial filtering to enhance filament detection. Quantitative analysis included filopodia number and actin fiber length using automated detection plugins, as described previously ^42^.

### Western Blot Analysis

Equal amounts of protein (30 µg) were resolved by SDS–PAGE and transferred onto nitrocellulose membranes. Membranes were blocked and incubated with primary antibodies targeting signaling proteins of interest (Table S1), followed by species-appropriate horseradish peroxidase–conjugated secondary antibodies. Immunoreactive bands were detected using enhanced chemiluminescence and imaged under non-saturating conditions.

Band intensities were quantified by densitometry using ImageJ v 5.5 software (NIH, USA) and normalized to total protein levels or housekeeping controls, as indicated for each experiment.

### Anti-TSP-1 neutralizing antibody treatment

To assess whether TSP-1 mediated the effects observed on brain endothelial cells induced by PE-CM, a neutralizing treatment was performed. PE-CM was preincubated with an anti-TSP-1 monoclonal antibody (Thermo Fisher Scientific, 1:500; v/v) overnight at 4°C. As a negative control, the CM was incubated in parallel with an isotype control anti-mouse IgG antibody (Thermo Fisher Scientific, 1:500; v/v). Subsequently, these pretreated media were used to stimulate hCMEC/D3 as described above for functional assays.

### TSP-1 Immunoprecipitation

To deplete TSP-1 from CM, immunoprecipitation was performed using Protein A/G Agarose Beads (Thermo Fisher Scientific, Waltham, MA, USA). CM (500 µg total protein) was incubated overnight at 4°C with an anti–TSP-1 antibody under gentle agitation. Protein A/G beads were then added and the mixture incubated for an additional 2 h at 4°C to allow immune complex capture. Beads were washed in ice-cold buffer (PBS containing 0.1% Triton X-100 and protease inhibitors) to remove unbound proteins. Bound proteins were eluted in reducing Laemmli buffer by heating at 100°C for 10 min. Eluates and corresponding input fractions were analyzed by Western blot to confirm TSP-1 removal.

### Proteomic analysis

Plasma samples were extracted from the umbilical cord, corresponding to women with normal pregnancy and women with preeclampsia (n=3 per condition), using a standardized protocol. Serums were depleted with the Multiple Affinity Removal Spin Cartridge Human-14 System, 0.45 mL. Then, in an SDS-PAGE gel, direct fractions (non-depleted serum), depleted fractions (direct flow) and eluted fractions (cartridge-bound proteins - more abundant) were analyzed. Each sample was prepared for mass spectrometry (nLC-MS/MS). 200 ng of the peptides obtained in the previous step were injected into a nanoElute nanoUHPLC (Bruker Daltonics, rapifleX®) coupled to the timsTOF Pro mass spectrometer (Trapped Ion Mobility Spectrometry – Quadrupole Time-of-Flight, Bruker Daltonics, Billerica, MA, USA).

### In silico analysis and bioinformatics

The resulting data were analyzed using the Dia-NN software with a spectral library of human serum (2,200 total proteins, developed by the MELISA Institute). Regarding the quantification of coefficient of linkage (CFL), the Perseus v1.6.15 software was used with the intensity values of each protein identified. Using the protein identification results, columns corresponding to Uniprot access codes and intensity values were selected. The data were concatenated and the missing values were entered in the intensity columns. The resulting matrix was normalized using Z-scores and medians. Differential expression proteins were determined by applying Student’s T-test with a Benjamini-Hochberg multiple correction test (p-value < 0.05). The identification of DEPs (Differentially Expressed Proteins) was carried out from a PE/NP comparison. For bioinformatic analyses, where protein-protein interactions were classified and predicted, version 1.2 (http://pantherdb.org/) of the PANTHER™ (Protein Analysis Through Evolutionary Relationships) classification system was used, together with the STRING v10 interaction network (http://www.string-db.org/) tool for the search and retrial of interacting genes/proteins (Table S2).

Proteins differentially expressed in preeclampsia were subjected to functional enrichment analysis based on Gene Ontology (GO) to identify significantly represented biological processes (p < 0.05). Subsequently, a hierarchical functional network was constructed using network analysis tools (Cytoscape), where “Preeclampsia” was defined as a central node, connecting with enriched functional clusters and their respective associated proteins.

The visualization made it possible to group related biological processes and evaluate the connectivity of proteins within the network. THBS1 was identified within the cluster associated with angiogenesis inhibition, highlighting its functional relevance in the pathophysiological context.

### Statistical Analysis

Data are presented as mean ± standard deviation (SD). Normality was assessed using the Shapiro–Wilk test. Depending on data distribution, comparisons between two groups were performed using unpaired Student’s t-test or Mann–Whitney test. For paired analyses (e.g., matched *in vitro* or tissue comparisons), paired t-tests were used. Multiple group comparisons were analyzed by one-way ANOVA followed by Dunn’s *post hoc* test. The unit of analysis (n) refers to independent biological replicates as specified in each figure legend. A two-tailed P value <0.05 was considered statistically significant. Statistical analyses were performed using GraphPad Prism software 8.0.1 version.

#### Sample size

Although a total of samples were available, not all functional assays were performed using the full cohort due to limitations in sample availability, primary cell viability after isolation, and technical constraints associated with mechanistic experiments. The number of biological replicates included in each experiment was determined based on sample availability and supported by sample size estimation considering an expected large effect size (Cohen’s d ≥ 1.0), an alpha level of 0.05, and a statistical power of 80%. Under these assumptions, a minimum of 5–7 biological replicates per group was considered sufficient to detect statistically significant differences.

## RESULTS

### Maternal and neonatal clinical characteristics

Considering diagnosis criteria and matched parameter, women who developed preeclampsia (n=26), delivered significantly earlier, and showed higher systolic and diastolic blood pressure than women with normal pregnancy (n=25; Table 1). Women with preeclampsia showed significant increase in proteinuria levels (0.75 ± 0.65 mg/mg vs 0.27 ± 0.3 mg/mg; p < 0.001). No statistically significant differences were found in parameters such as maternal age, parity, mode of delivery, or neonatal antropometry or placental weight.

**Table 1.** Maternal and neonatal clinical characteristics of the studied groups.

| Parameter | Normal pregnancy<br>(n=25) | Preeclampsia<br>(n=26) | p value |
| --- | --- | --- | --- |
| *Maternal characteristics* |  |  |  |
| Maternal age (years) | 26.8 ± 5.2 | 29.4 ± 4.8 | 0.08 |
| Parity. n (%) |  |  |  |
| Primiparus | 16 (64%) | 12 (46%) | 0.31 |
| Multiparus | 9 (36%) | 14 (54%) |  |
| Gestational age at delivery (weeks) | 39.2 ± 1.1 | 37 ± 2.5 | <0.001 |
| SBP (mmHg) | 118.3 ± 5.6 | 142.6 ± 18.2 | <0.001 |
| DBP (mmHg) | 68.4 ± 9.3 | 90.7 ± 13.8 | <0.001 |
| Ratio (Prot/Cr) | 0.3 ± 0.3 | 0.7 ± 0.6 | <0.001 |
| Mode of delivery |  |  |  |
| Vaginal | 18 (72%) | 15 (57.6 %) | 0.43 |
| Cesarean | 7 (28%) | 11 (42.4) |  |
| *Neonatal characteristics* |  |  |  |
| Sex. n (%) |  |  |  |
| Male | 12 (48%) | 15 (58%) | 0.68 |
| Female | 13 (52%) | 11 (42%) |  |
| Birth weight (g) | 3119 ± 989 | 2322 ± 1480 | 0.07 |
| Newborn size (cm) | 49.3 ± 1.4 | 61.9 ± 72.6 | 0.21 |
| Placenta weight (g) | 620.6 ± 180.2 | 864.6 ± 1252 | 0.80 |

SBP, Systolic blood pressure. DBP, Diastolic blood pressure. HUVEC: human umbilical vein endothelial cells; g: grams; cm: centimeters; Prot: proteins; Cr: creatinine

### Preeclampsia-derived endothelial secretome impairs angiogenic capacity of human brain endothelial cells

To determine whether the fetoplacental endothelial secretome influences brain endothelial function, human brain microvascular endothelial cells (hCMEC/D3) were exposed to NP-CM or PE-CM (Figure 1A).

**Figure 1.**
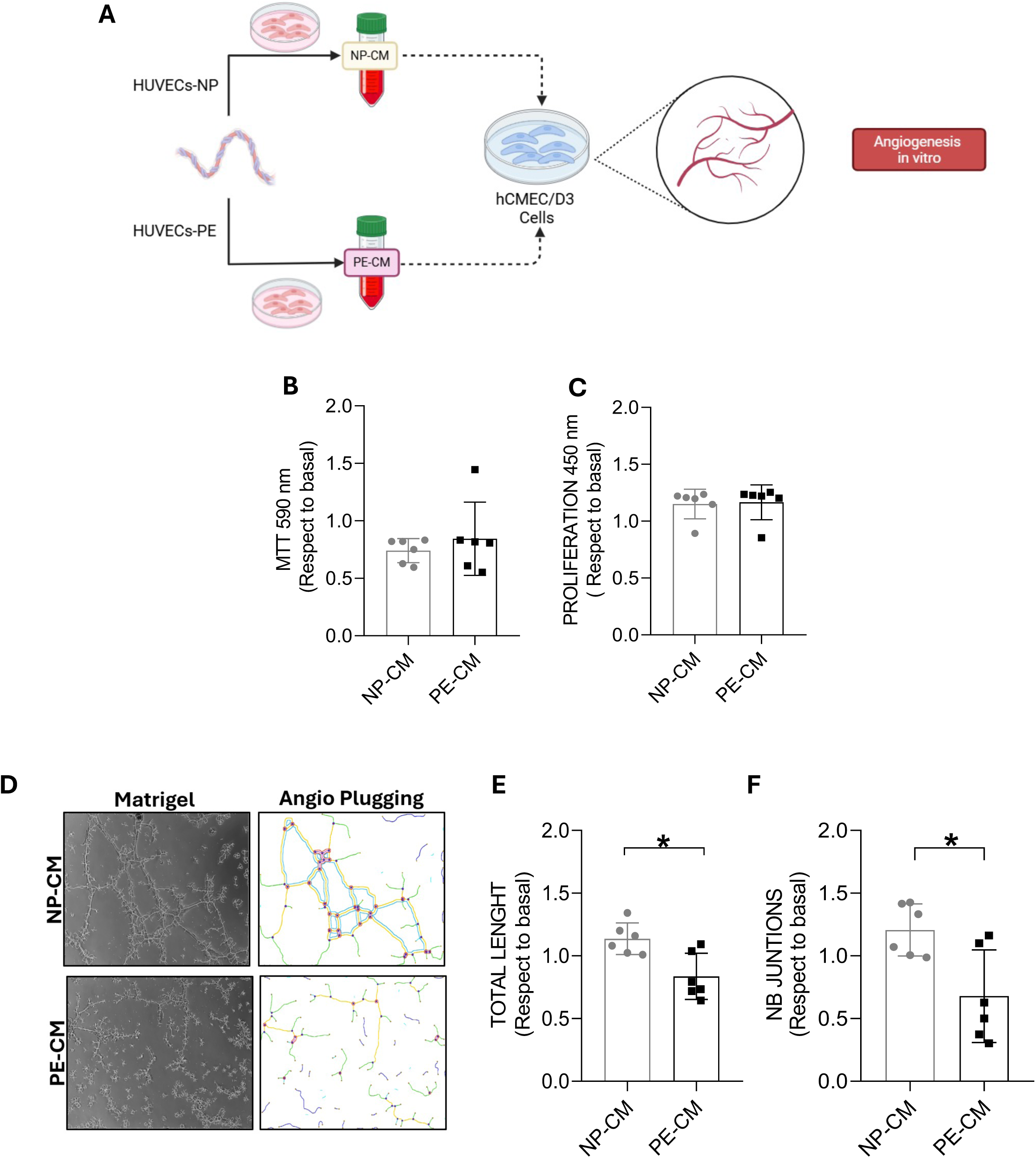
The conditioned medium from PE-HUVECs reduce angiogenic capacity of brain endothelium. **(A)** Experimental design scheme. hCMEC/D3 cells were treated with conditioned medium (CM) derived from normal pregnancy HUVECs (NP-CM) or preeclampsia (PE-CM) and then angiogenesis *in vitro* was evaluated. **(B)** Metabolic activity assay (MTS) in hCMEC/D3 treated (6 h) with NP-CM or PE-CM. Data presented as a fold of change to control (i.e., without treatment)(n=6 per condition). **(C)** Proliferation assay in hCMEC/D3 after (24 h) of treatment with NP-CM or PE-CM (n=6 per condition). **(D)** Representative images of the angiogenesis *in vitro* assay in hCMEC/D3 cells stimulated with the NP-CM and PE-CM. **(E)** Quantification of the total length of tubes. And **(F)** number of junctions, normalized with respect to the control (n=6 per condition). Each dot represents one biological replicate corresponding to CM derived from an individual patient. Data expressed as mean ± SD. Statistical analysis using Mann-Whitney test. *p<0.05.

Exposure to PE-CM for 6 h did not alter hCMEC/D3 viability or proliferation compared with NP-CM (Figure 1B–C), indicating that subsequent functional effects were not attributable to cytotoxicity or growth suppression. In contrast, PE-CM markedly impaired the angiogenic capacity of brain endothelial cells *in vitro* (Figure 1D-1F). Angiogenesis *in vitro* analysis revealed significant reductions in total tube length (Figure 1E), number of junctions (Figure 1F), and branch points (Figure S1A) following PE-CM treatment compared with NP-CM.

These findings indicate that the umbilical endothelial secretome from preeclampsia suppresses angiogenic function in brain endothelial cells without affecting cell survival.

### Preeclampsia endothelial secretome suppresses migration and invasive behavior of brain endothelial cells

Because endothelial migration is essential for brain angiogenesis ^43^ we examined whether PE-CM alters motility of human brain endothelial cells. Time-course analysis revealed that NP-CM stimulated hCMEC/D3 migration, with maximal responses between 6–9 h. In contrast, PE-CM failed to induce a comparable migratory response and resulted in significantly reduced cell movement during early time points (Figure 2A–B). Integrated analysis using area under the curve (AUC) confirmed an overall suppression of endothelial migration under PE-CM compared with NP-CM (NP-CM: 41.06 ± 7.93 vs. PE-CM: 28.50 ± 2.03 AUC, p= 0.0469).

**Figure 2.**
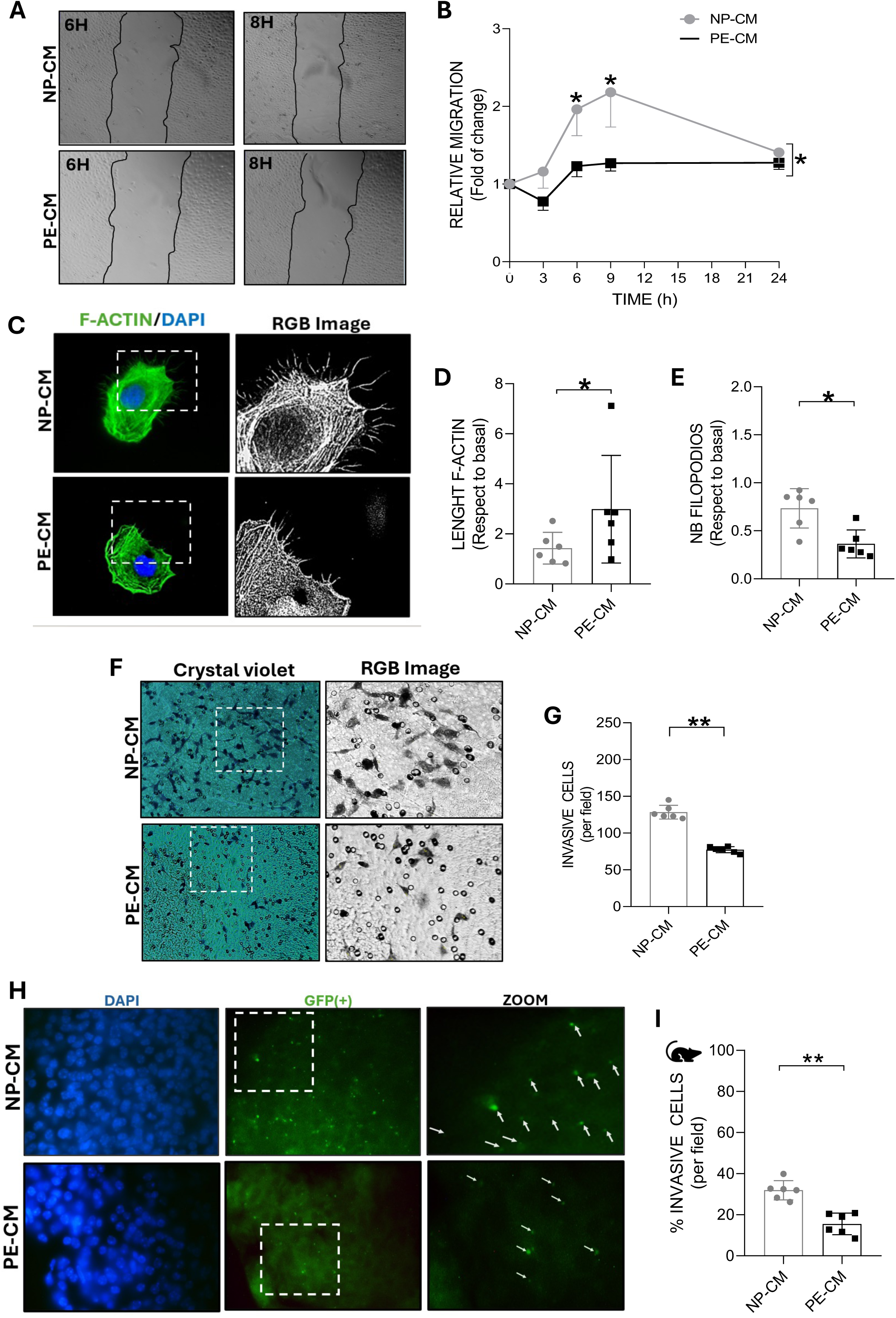
The conditioned medium from PE-HUVEC reduce cell migration/invassion in brain endothelial cells. **(A)** Representative images of the wound healing assay (3h to 24h) to measure the migratory capacity of hCMEC/D3 cells treated with umbilical endothelial conditioned medium (NP-CM or PE-CM). **(B)** Relative migration area normalized to control condition (i.e., without treatment)(n = 6 per condition) **(C)** Representative images of F-actin in hCMEC/D3 cells treated with NP-CM or PE-CM. **(D)** Quantification of the length of actin filaments for the total area in cells treated as C. Values presented with respect to control (n=6 per condition). **(E)** Number of filopodia in cells treated as in C. **(F)** Representative images of the cell invasion assay observed in cells hCMEC/D3 treated with NP-CM and PE-CM. **(G)** Number of invasive cells per field (n=6 per condition). **(H)** Representative images of brain slices cultured in Transwell® inserts and incubated (24 h) with GFP-labeled human brain endothelial cells (hCMEC/D3-GFP). NP-CM or PE-CM was used as chemoattracting factor. Boxed areas are presented at higher magnification. Arrows denote hCMEC/D3-GFP–positive cells within the brain parenchyma. **(I)** GFP+ cells/DAPI+ cells per section. Each dot represents one biological replicate corresponding to CM derived from an individual patient. Data expressed as mean ± SD. Statistical analysis using Mann-Whitney test and Paired T test *p<0.05, **p<0.01.

Given the dependence of migration on cytoskeletal dynamics, F-actin organization was assessed. PE-CM exposure altered actin architecture, characterized by increased actin fiber length and a reduction in filopodia formation relative to NP-CM–treated cells (Figure 2C–E), indicating impaired cytoskeletal remodeling.

We next evaluated brain endothelial invasion, a process requiring coordinated migration and matrix interaction. Validation of our cell invasion assay *in vitro* is presented in Figure S1B. We found that PE-CM significantly reduced the invasiveness capacity of hCMEC/D3 cells *in vitro* compared with NP-CM (Figure 2F–G), consistent with a motility-restricted phenotype.

To test whether this anti-migratory effect persisted in a complex tissue environment, GFP-labeled hCMEC/D3 cells were applied to organotypic brain slice cultures (Figure S1C-S1D). Our results indicate that after 24 h, NP-CM–treated cells infiltrated multiple tissue planes, whereas PE-CM–treated cells remained predominantly at superficial layers. Quantification demonstrated a significant reduction in the number of invading endothelial cells under PE-CM conditions (Figure 2H–I).

Together, these findings show that the umbilical endothelial secretome from preeclampsia disrupts cytoskeletal organization and suppresses both migratory and invasive processese in brain endothelial cells.

### Functional validation of PE-CM effects on fetal brain endothelium

Because hCMEC/D3 cells represent adult endothelium, we next determined whether PE-CM–induced dysfunction is also present in fetal brain endothelial cells. Primary brain microvascular endothelial cells (BMECs) were isolated from E18.5 embryos of normotensive dams (Sham) and RUPP dams (PE-like model). To distinguish CM-driven effects from intrinsic endothelial susceptibility, BMECs from both origins were exposed to NP-CM or PE-CM, and angiogenic capacity was assessed *in vitro* (Figure 3A).

**Figure 3.**
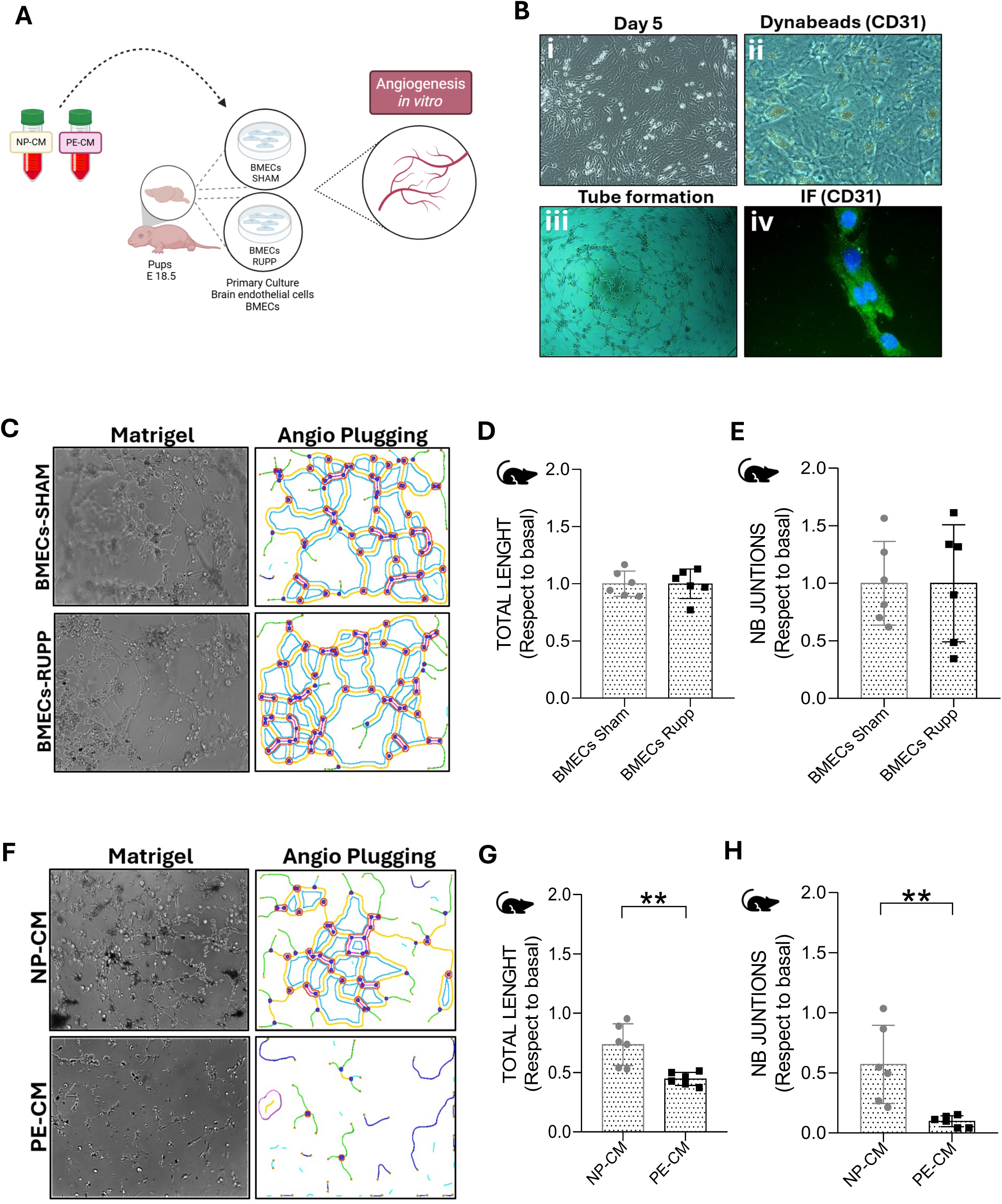
PE-CM reduce angiogenic capacity of fetal brain endothelial cells of PE-like conditions. **(A)** Experimental model used. Microvascular fetal brain endothelial cells (BMECs) from SHAM or RUPP fetuses (murine model of preeclampsia) were exposed to NP-CM or PE-CM, respectively to evaluate *in vitro* angiogenesis. **(B)** Characterization of BMEC (i) Day 5 post *in vitro* culture. (ii) Immunoselection of endothelial cells with CD31+ coated beads. (iii) *In vitro* angiogenesis in control condition (6h). (iv) Immunofluorescence of CD31 (green) and nucleus (DAPI). **(C)** Representative images of *in vitro* angiogenesis in BMECs from SHAM and RUPP pups under basal conditions (without any stimulus). **(D)** Quantification of total tube length, and **(E)** number of junctions (n=6 per condition). **(F)** BMECs isolated from SHAM offspring were exposed to NP-CM, whereas BMECs from RUPP offspring were exposed to PE-CM. **(G)** Quantification of the total length, and **(H)** number of junctions (n=5 per condition). Each dot represents one biological replicate corresponding to CM derived from an individual patient. Data expressed as mean ± SD. Statistical analysis using Mann-Whitney test **p<0.01

We first confirmed the endothelial identity of isolated BMECs (Figure 3B), identifying typical endothelial morphology in culture, attachment of CD31-magnetic beads for immunoselection, formation of capillary-like networks in Matrigel, and CD31 immunoreactivity in the CD31 positively immunoselected cells.

We then evaluated baseline angiogenic capacity in BMECs from Sham and RUPP embryos (Figure 3C). Under control conditions, no differences were observed between groups, as indicated by comparable total tube length (Figure 3D), number of junctions (Figure 3E), and number of branches (Figure S1E).

In contrast, exposure to PE-CM revealed a selective vulnerability of RUPP-derived BMECs. Thus, compared with NP-CM, PE-CM significantly reduced angiogenic capacity in BMECs from RUPP embryos, reflected by decreased total tube length (Figure 3F-G), fewer junctions (Figure 3H), and reduced branching (Figure S1F).

These findings indicate that umbilical endothelial secretome from preeclampsia impair fetal brain endothelial angiogenesis, an effect that is amplified in endothelium originating from a PE-like intrauterine environment.

### TSP-1 is increased in the fetal environment in preeclampsia

To identify fetal circulating mediators that could explain the anti-angiogenic effects observed in our models, we performed comparative proteomic profiling of umbilical cord serum from normal pregnancies (n=3) and preeclampsia (n=3). Quantitative mass spectrometry identified 984 proteins, of which 30 were differentially expressed proteins (DEPs) in samples from preeclamptic pregnancies relative to those from normal pregnancy (Table S2). Among these, 22 proteins were upregulated and 8 were downregulated in preeclampsia (Figure 4A, Figure S2).

**Figure 4.**
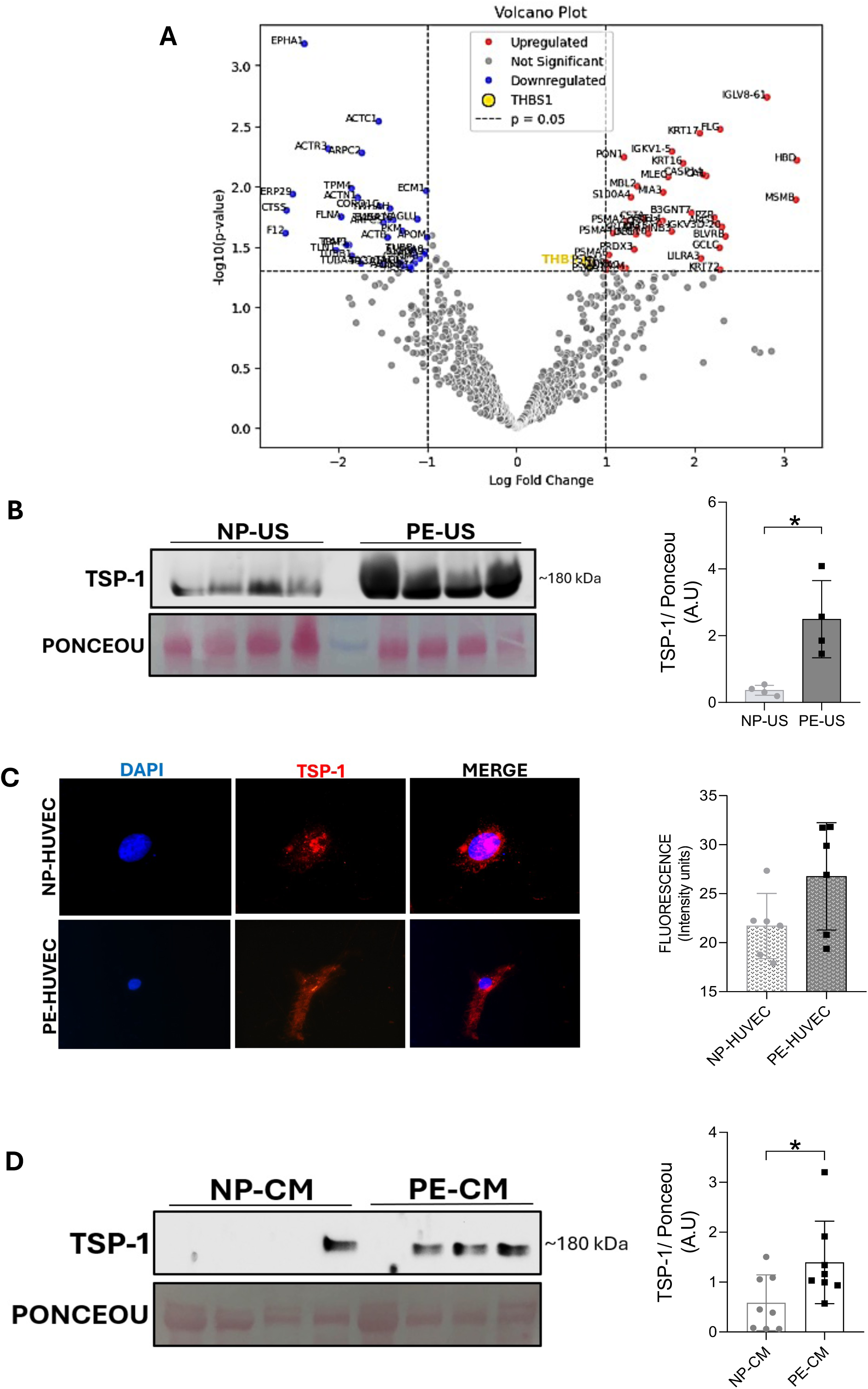
Increased thrombospondin 1 in the fetoplacental setting of PE. **(A)** Volcano plot of the proteomic analysis of umbilical cord serum of women with normal pregnancy (NP) and preeclamptic pregnancy (PE) (n=3 per condition). Upregulated (red color) and downregulated (blue color) are remarked. Thrombospondin 1 (THBS1 or TSP-1) is highlited in yellow. **(B)** Western blot analysis of TSP-1 on individual samples of umbilical cord serum from NP and PE pregnancies (n=4 per condition). **(C)** Immunofluorescence for TSP-1 in NP-HUVEC and PE-HUVEC. TSP-1 estimation in fluorescence intensity as showed in C. **(D)** TSP-1 expression in NP-CM and PE-CM. Ponceau staining was used as loading control (n=8 per condition). Each dot represents one biological replicate corresponding to CM derived from an individual patient. Data expressed as mean ± SD. Statistical analysis using Mann Whitney test *p<0.05.

Gene Ontology (Biological Process) enrichment analysis of upregulated proteins in umbilical plasma of preeclamptic pregnancies revealed overrepresentation of pathways related to gas transport (including oxygen and carbon dioxide), hydrogen peroxide metabolism, and oxidative stress–related processes, suggesting altered redox homeostasis in the fetal compartment (Figure S3A). Enrichment was also observed in processes associated with intermediate filament cytoskeleton organization, peptide crosslinking, and long-chain fatty acid transport, consistent with structural and metabolic remodeling of the fetoplacental vascular environment.

Among the upregulated candidates, TSP-1, a well-established endogenous inhibitor of angiogenesis, showed a 1.5-fold increase in umbilical plasma of preeclamptic pregnancies (Figure 4A) compared to normal pregnancy. This elevation was independently confirmed by Western blot in additional samples (n=4 per group; Figure 4B).

Pathway analysis using STRING further supported the biological relevance of TSP-1, placing it within networks regulating angiogenesis inhibition, extracellular matrix remodeling, cell adhesion, and chemotaxis (Figure S3B). Likewise, the analysis of protein-protein interaction of TSP-1 by STRING showed a network composed of 19 nodes and 19 interactions, with an average degree of connectivity of 2 and a local clustering coefficient of 0.565 (Figure S3C).

To determine whether endothelial cells contribute to this TSP-1–enriched environment, we assessed its expression and secretion in HUVECs derived from normal (n=6) and preeclamptic (n=6) pregnancies. Cellular TSP-1 expression did not differ between groups (Figure 4C). However, TSP-1 levels were significantly higher in PE-CM compared with NP-CM (Figure 4D), indicating enhanced extracellular release rather than altered intracellular abundance. In contrast, secretion of another anti-angiogenic mediators commonly linked to preeclampsia, the decoy receptor of VEGF, sFlt-1 (Figure S4A), was not different between PE-CM and NP-CM.

Together, these findings identify TSP-1 as a prominent anti-angiogenic factor enriched in the fetal circulation in preeclampsia and support its role as a candidate mediator of the PE-CM antiangiogenic effect observed in brain endothelial cells.

### TSP-1 interacts with VEGF and modulates VEGFR2-dependent angiogenic signaling

Given the elevated levels of TSP-1 in PE-CM and its established anti-angiogenic role, we investigated whether TSP-1 interferes with VEGF-driven pro-angiogenic signaling (Figure S5A).

VEGF levels were first assessed in NP-CM and PE-CM. Using Western blot, VEGF was not detectable neither in NP-CM nor in PE-CM under regular experimental conditions (Figure 5A), despite detection of the positive control, a recombinant VEGF. Ponceau staining verified comparable protein loading across those analyzed samples (Figure S5B).

**Figure 5.**
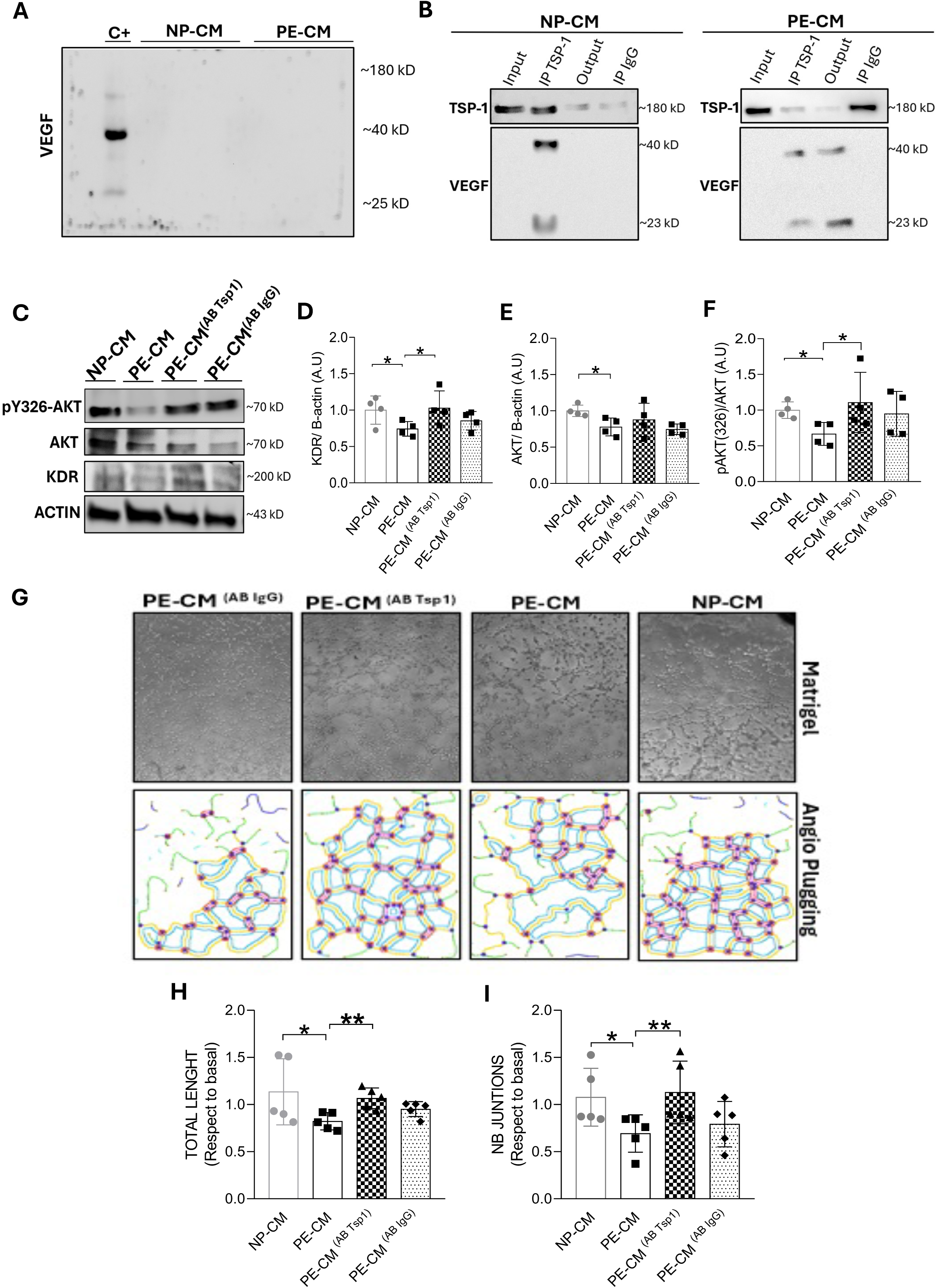
TSP-1 sequester VEGF in PE-CM and impairs VEGFR2 downstream signaling pathway. **(A)** Representative Western blot showing VEGF detection using placental tissue lysate as a positive control (C+). No VEGF signal was detected in NP-CM or PE-CM samples (n = 3 per condition). **(B)** TSP-1 immunoprecipitation (IP) in NP-CM and PE-CM samples. TSP-1 and VEGF were western blotting. Input (total CM), Output (remant after IP), and Negative Control (IP IgG) controls are included. **(C)** Represenative Western blot of hCMEC/D3 treated (6 h) with NP-CM, PE-CM, PE-CM + neutralizing antibody against TSP-1 (PE-CM ^AB^ ^TSP1^), and PE + IgG control antibody (PE-CM ^AB^ ^IgG^). **(D)** Densitometry of KDR, **(E)** AKT, and **(F)** pAKT (325) protein levels as in C (n=4 per condition). β-Actin was used as loading control. **(G)** Representative images *in vitro* angiogenesis in hCMEC/D3 treated as in C. **(H)** Total length of the tubes and **(I)** Number of junctions of the assay shown in G (n=5 per condition). Each dot represents one biological replicate corresponding to CM derived from an individual patient. Data expressed as mean ± SD. Statistical analysis using ANOVA followed by post hoc Kruskal Wallis test. *p<0.05 **p<0.01.

Then, we hypothetize that TSP-1 can sequester VEGF reducing its detection and biological availability. To test this, we used CM to immunoprecipitated TSP-1 and then followed by VEGF detection by Western blot. VEGF was detected in TSP-1 immunoprecipitates from both NP-CM and PE-CM, suggesting a physical association between these two proteins (Figure 5B). Notably, analysis of the post-immunoprecipitation supernatant revealed reappearance of detectable VEGF only in PE-CM, but not in NP-CM. This finding is consistent with a greater fraction of VEGF being retained within TSP-1–containing complexes in the preeclamptic condition.

We next examined the impact of this interaction on VEGFR2 signaling in the brain endothelial cell line (hCMEC/D3) (Figure 5C). Exposure to PE-CM significantly reduced total VEGFR2 protein levels (Figure 5D), AKT abundance (Figure 5E), and AKT phosphorylation (Figure 5F) compared with NP-CM. To confirm TSP-1 participation, we found that TSP-1 antibody neutralization restored VEGFR2 expression and AKT phosphorylation. Confirming that TSP-1 is a key mediator of PE-CM–induced suppression of VEGF/VEGFR2 signaling (Figure 5D, 5F).

We then assessed whether TSP-1 blockade could functionally rescue angiogenesis. As previously observed, PE-CM markedly impaired angiogenesis *in vitro* in hCMEC/D3 cells (Figure 5G). TSP-1 antibody-mediated neutralization significantly restored angiogenic capacity, normalizing total tube length (Figure 5H) and junction number (Figure 5I) to levels comparable to NP-CM–treated cells. Consistently, PE-CM depleted of TSP-1 by immunoprecipitation showed reduced anti-angiogenic activity (Figure S5C–E).

Together, these results identify TSP-1 as a functional mediator of PE-CM–induced anti-angiogenic effects, acting through sequestration of VEGF and suppression of VEGFR2–AKT signaling in brain endothelial cells.

## DISCUSSION

We uncover a previously unrecognized form of linking the fetoplacental and brain vasculatures in preeclampsia. The umbilical vein endothelial cell secretome imposes an antiangiogenic phenotype on brain endothelial cells, impairing migration, invasion, and network formation. Mechanistically, TSP-1 emerges as a key mediator. TSP-1 is enriched in the fetal environment, it binds VEGF, restricts VEGFR2–AKT signaling, and drives brain endothelial dysfunction, while its neutralization restores angiogenic capacity. These findings identify disrupted vascular crosstalk as a mechanism by which preeclampsia may limit fetal brain angiogenesis, potentially programming acute and long-term neurovascular vulnerability in the offspring.

There is increasing evidence that preeclampsia exerts a profound and lasting impact on the development of the offspring’s central nervous system, ^44–48^ particularly through alterations in the formation, maturation, and functionality of the cerebral vasculature ^11,13,49^. Epidemiological studies indicate that offspring of women with preeclampsia exhibit an increased risk of neurovascular complications, including perinatal stroke, cerebral edema, and hemorrhagic events ^5^, although the underlying mechanisms remain poorly defined. Our data provide a mechanistic link between the adverse fetoplacental environment and impaired cerebrovascular development. We previously demonstrated reduced brain angiogenesis ^11^ and increased blood–brain barrier permeability ^13^ in preclinical models of preeclampsia. These findings align with clinical evidence of altered cerebral perfusion in children born to women with preeclampsia ^50^ and experimental reports of impaired cerebrovascular autoregulation in related models ^51^. Current findings support the concept that early endothelial–endothelial dysregulation in preeclampsia may program a vulnerable cerebrovascular phenotype in the offspring. Furhtermore, these cerebrovascular complications may compromise the cognitive, behavioral, and neurodevelopment later in life^11,13,52,53^.

Our data support that fetoplacental-derived circulating factors, rather than intrinsic endothelial defects, drive impaired brain angiogenesis in preeclampsia. Primary brain endothelial cells from RUPP offspring preserved baseline angiogenic capacity, but exhibited reduced angiogenesis *in vitro* when exposed to PE-CM, indicating an environmentally induced antiangiogenic phenotype. These findings integrate with prior reports of reduced cerebral angiogenesis ^11^, impaired neonatal perfusion ^12^ and suppressed KDR signaling in the RUPP model ^54,55^, positioning circulating placental mediators as key regulators of altered neurovascular development in preeclampsia.

Our results support previous findings showing circulating TSP-1 in umbilical vein serum, which respond to changes in the fetal compartment ^56^. Thus, circulating fetal TSP-1 levels negatively correlated with fetal venous pH, but positively correlated with fetal VEGF levels. In addition, the authors reported a significant increase in fetal TSP-1 levels in cases of intrauterine growth restriction, positioning this glycoprotein as a likely sensitive marker of the fetal vascular environment.

TSP-1 is classically recognized as an antiangiogenic factor that acts directly through VEGF sequestration and inhibition of VEGFR2 (KDR) activation ^57–59^ Moreover, TSP-1 can also inhibit VEGF signaling through interactions with receptors such as CD36 and CD47, promoting antiangiogenic and pro-apoptotic effects in endothelial cells ^28,60–62^. Effects that involve suppression of key intracellular cascades such as PI3K/AKT/eNOS ^63–65^. Our results extend this framework to fetoplacental–brain crosstalk in preeclampsia. TSP-1 was enriched in the fetal endothelial secretome and umbilical plasma, associates with VEGF, reduces VEGFR2/AKT signaling in brain endothelial cells, and drives antiangiogenic dysfunction. An effect that was prevented with TSP-1 blockade, showing the critical relevance of this factor in the fetal endothelium secretoma of preeclamptic pregnancies.

The origin of TSP-1 in umbilical cord plasma is likely fetal endothelial cells, such as HUVEC, as these cells express and release TSP-1, as shown in NP-CM and PE-CM. Moreover, the release of TSP-1 was higher in preeclampsia than in normal pregnancy-derived HUVECs. These results complement other findings showing TSP-1 synthesis predominantly in the placental compartment, particularly the syncytiotrophoblast ^30,66,67^. In this regard, higher TSP-1 circulating or placental levels has been previously analyzed in the context of maternal endothelial dysfunction present in women with preeclampsia ^29,31^. However, TSP-1 maternal levels were decreased in severe preeclampsia (i.e, HELLP syndrome), suggesting a progressive dysregulation of the angiogenic system in advanced stages of the disease ^31,32^. No previous studies have analyzed circulating TSP-1 levels in the umbilical cord in the context of preeclampsia.

Endothelial-endothelial communication represents another central aspect of this work. The endothelium is recognized as an active paracrine and endocrine organ capable of releasing soluble proteins, extracellular vesicles, microRNAs, and angiogenic factors that act as autocrine and paracrine molecules ^68,69^. In this framework, the placenta–fetal brain axis has emerged as a central focus for understanding the mechanisms underlying the alterations observed in offspring of mothers with pathological pregnancies, including preeclampsia ^70,71^. Several studies have shown that placental-derived antiangiogenic factors, such as sFlt-1, are upregulated in the fetal circulation and are associated with reduced angiogenesis in fetal endothelial cells ^72,73^. More recently, it has been reported that placental insulin-like growth factor 1 (IGF1) overexpression leads to altered placental function in a sex-specific manner, and these defects were associated with brain development (i.e., striatal development, a functional region within the forebrain that contributes to motor regulation and procedural and habit learning). Since IGF1 is primarily expressed in the endothelial cells of the fetal region of the human placenta and the labyrinth zone of the mouse placenta ^74,75^ these results connect directly alterations in vascular placental and brain development. Building on these results, our study uncovers an endothelial-endothelial communication during impaired brain angiogenesis in offspring of preeclampsia.

Our proteomic analysis did not detect significant sFlt-1 overexpression in fetal plasma, reinforcing TSP-1 as an alternative, and potentially dominant, negative regulator of fetal angiogenesis. Our immunoprecipitation assays showed evidence of physical association between TSP-1 and VEGF in PE-CM. Functionally, TSP-1–enriched PE-CM reduced total VEGFR2 abundance in brain endothelial cells and suppressed downstream AKT activation, indicating inhibition of the VEGFR2/AKT axis. This signaling disruption was mechanistically linked to impaired endothelial angiogenesis and was fully prevented by TSP-1 neutralization, confirming its central role in the antiangiogenic effects of PE-CM.

This study has limitations. Direct *in vivo* validation of TSP-1–dependent cerebrovascular remodeling will be required to establish the physiological magnitude and temporal dynamics of this pathway in the developing brain. In addition, while our data support functional interference with VEGF/VEGFR2 signaling, the precise molecular mechanism (whether through ligand sequestration, receptor internalization, or modulation of receptor stability), remains to be defined. Nevertheless, the convergence of unbiased proteomics with orthogonal functional and neutralization approaches across independent endothelial systems provides strong mechanistic evidence that TSP-1 is a critical regulator of impaired fetal cerebral angiogenesis in preeclampsia.

In conclusion, TSP-1 emerges as a central mediator of the antiangiogenic phenotype imposed on cerebral endothelium by the fetal endothelial secretome in preeclampsia. By sequestering VEGF and suppressing VEGFR2–AKT signaling, TSP-1 limits brain endothelial angiogenesis. These findings uncover a previously unrecognized endothelium-to-endothelium signaling axis linking the fetoplacental compartment to the cerebral vasculature, and provide a mechanistic framework for how preeclampsia may program acute cerebrovascular vulnerability in the offspring.

## ACKNOWLEDGMENT

The authors thank the Vascular Physiology Laboratory and GRIVAS Health researchers for their valuable input.

## SOURCE OF FUNDING

This study was funded by Fondecyt 1240295 (Chile). HDM was funded by a British Heart Foundation Intermediate Basic Science Fellowship (FS/15/32/31604).

## DISCLOSURE

The authors declare that they have no conflict of interest.

## DATA AVAILABILITY

Data is available upon reasonable request to Prof. Carlos Escudero.

## AUTHOR CONTRIBUTION

CE conceptualized the manuscript. EGG performed most of the experiments and analyses. JA, HS, FT performed some animal and *in vitro* experiments. CV carried out the proteomic analysis. MA performed bioinformatics studies of protein interaction. HDM and LOK were in charge of patient selection for umbilical cord analysis. RMC was advisor in analysis of the conditioned medium and experimental approach. All co-authors approved the final version of this manuscript.

## ARTIFICIAL INTELLIGENCE USE

Artificial intelligence (ChatGPT, OpenAI) was used exclusively to improve the clarity and grammar of the manuscript. No data were generated, analyzed, or interpreted using artificial intelligence. The authors take full responsibility for the content of the manuscript.

## Abbreviations

PE: Preeclampsia
NP: normal pregnancy
CM: conditioned media
HUVECs: umbilical vein endothelial cells
HUVECs-NP: HUVECs from normal pregnancy
HUVECs-PE: HUVECs from preeclamptic pregnancy
NP-CM: conditioned media from HUVECs-NP
PE-CM: conditioned media from HUVECs-PE
BMECs: microvascular brain endothelial cells
hCMEC/D3: brain microvascular endothelial cell
TSP-1: Trombospondin 1
VEGF: vascular endothelial growth factor
VEGFR2/KDR: vascular endothelial growth factor receptor 2
pKDR: phospho-VEGF Receptor 2 (Tyr951)
RUPP: reduced uterine perfusion pressure
PlGF: placental growth factor
PBS: Phosphate-Buffered Saline
FBS: Fetal bovine serum
BSA: Bovine Serum Albumin
PFA: paraformaldehyde
sFLT-1: soluble Fms-like tyrosine kinase-1
PI3K: Phosphoinositide 3-kinase
eNOS: Endothelial Nitric Oxide Synthase
HELLP Syndrome: hemolysis elevated liver enzymes and low platelet count syndrome
IGF1: insulin-like growth factor 1

## Notes

**Conflict of interest statement:** The authors declared no conflict of interest.

### Competing Interest Statement

The authors have declared no competing interest.

